# Pumilio protects Xbp1 mRNA from regulated Ire1-dependent decay

**DOI:** 10.1101/2021.02.08.430300

**Authors:** Fatima Cairrao, Cristiana C Santos, Adrien Le Thomas, Scot Marsters, Avi Ashkenazi, Pedro M. Domingos

## Abstract

The unfolded protein response (UPR) maintains homeostasis of the endoplasmic reticulum(ER). Residing in the ER membrane, the UPR mediator Ire1 deploys its cytoplasmic kinase-endoribonuclease domain to activate the key UPR transcription factor Xbp1 through non-conventional splicing of Xbp1 mRNA. Ire1 also degrades diverse ER-targeted mRNAs through regulated Ire1-dependent decay (RIDD), but how it spares Xbp1 mRNA from this decay is unknown. We identified binding sites for the RNA-binding protein Pumilio in the 3’UTR *Drosophila* Xbp1. In the developing *Drosophila eye*, Pumilio bound both the Xbp1^unspliced^ and Xbp1^spliced^ mRNAs, but only Xbp1^spliced^ was stabilized by Pumilio. Furthermore, Pumilio displayed Ire1 kinase-dependent phosphorylation during ER stress, which was required for its stabilization of Xbp1^spliced^. Human IRE1 could directly phosphorylate Pumilio, and phosphorylated Pumilio protected Xbp1^spliced^ mRNA against RIDD. Thus, Ire1-mediated phosphorylation enables Pumilio to shield Xbp1^spliced^ from RIDD. These results uncover an important and unexpected regulatory link between an RNA-binding protein and the UPR.

## INTRODUCTION

Metazoan cells respond to endoplasmic reticulum (ER) stress by activating an intracellular network of signaling pathways, known as the unfolded protein response (UPR)^1,2^. In higher eukaryotes, the UPR involves three ER transmembrane transducers: inositol-requiring enzyme 1 (IRE1), pancreatic ER kinase (PKR)-like ER kinase (PERK), and activating transcription factor 6 (ATF6). When misfolded proteins accumulate in the ER, IRE1 activates the downstream transcription factor X-box binding protein 1 (XBP1), via the non-conventional splicing of Xbp1 mRNA^3–7^. The cytoplasmic kinase-endoribonuclease domain of IRE1 mediates the splicing of a 26-nucleotides long intron from the Xbp1 mRNA, causing a frame-shift during translation that introduces a new carboxyl domain in the XBP1 protein. The resulting spliced form of Xbp1, XBP1^spliced^, is a functionally active transcription factor that upregulates the expression of ER chaperones and other UPR target genes^8^.

The intracellular localization, stability and translation of mRNAs is regulated by interaction of RNA-binding proteins (RBPs) or microRNAs with specific sequences present in the 3’ untranslated regions (3’UTR) of the mRNA^9–11^. In budding yeast, the 3’UTR of the Xbp1-orthologue Hac1 targets Hac1 mRNA to foci of activated IRE1 at the ER membrane, which enables IRE1-mediated splicing of Hac1 mRNA^12^. In contrast, in mammalian cells the 3’UTR of Xbp1 seems to be dispensable for the targeting of Xbp1 mRNA to activated IRE1^13^. Instead, the targeting occurs through the tethering of 2 hydrophobic regions (HR1 and HR2) present in the XBP1^unspliced^ protein to the cytosolic side of the ER membrane^13,14^. This latter process involves a tripartite complex comprising Xbp1 mRNA, a ribosome, and the nascent XBP1 protein, and additionally requires translational pausing of the Xbp1^unspliced^ mRNA^14^.

Besides Xbp1 mRNA splicing, the endoribonuclease (RNase) domain of Ire1 also performs a function known as regulated IRE1-dependent decay (RIDD), which mediates the depletion of specific mRNAs^15^ and/or microRNAs^16^. While in *Drosophila* cells RIDD degrades ER-targeted mRNA relatively promiscuously, in mammalian cells it is thought to depend on a specific Xbp1-like mRNA sequence endomotif within a stem-loop structure, and on translational state of the mRNA target^17^. The phosphorylation and oligomerization state of IRE1 plays an additional role in controlling IRE1’s RNase activity^18^. RIDD can act as a post-transcriptional mechanism to deplete mRNAs, thereby affecting both ER homeostasis and cell fate^19,20,21,22^.

RNA-binding proteins (RBPs) constitute an important class of post-transcriptional regulators. RBPs are involved in multiple critical biological processes, relevant to cancer initiation, progression, and drug resistance^23^. Several recent studies validate the role of the Pumilio family of RBPs in diverse biological processes in several organisms. Specific mRNA targets of two human Pumilio isoforms (PUM1 and PUM2) include oncogenes, tumor suppressors, and other factors implicated in oncogenic and cell death pathways^24–26^. In several organisms, PUM proteins also play a conserved role in stem cell proliferation and self-renewal^27–30^. PUM proteins are essential for development and growth, and their dysfunction has been associated with neurological diseases, infertility, movement disorders, and cancer^31–36^.

*Drosophila* Pumilio - the founding member of the PUM family – Is characterized as a translational repressor and is involved in embryo patterning, fertility and the regulation of neuronal homeostasis^31,37–40^. PUM proteins act as post-transcriptional regulators, by interacting with consensus sequences called Pumilio regulatory elements (PRE) in the 3’UTRs of target mRNAs^9^ to modulate their translation and/or degradation^41–42^. Recent findings support a direct role of PUM proteins in the activation/protection of specific RNAs^26^. PUM proteins are composed of distinct functional domains: N-terminal repressor domains (N), which are unique to different PUM orthologues, and a Pumilio homology domain (PumHD), which recognizes the PRE. These domains mediate the normal repressive role of PUM proteins by antagonizing the translational activity of Poly(A) binding protein (PABP)^42,43^. The PUM N terminus is required to fully rescue developmental defects of a *pum* mutant^44,45^.

Although much progress has been made toward understanding the biological roles of PUM proteins, it remains to be determined how they regulate their target mRNAs particularly under diverse biological conditions such as cellular stress. Here, we uncover an unexpected functional link between Pumilio and the Ire1-Xbp1 pathway. We show that Pumilio protects *Drosophila* Xbp1 against RIDD, without perturbing canonical Ire1-driven Xbp1 splicing. Furthermore, we provide evidence that Ire1 phosphorylates Pumilio under ER stress and that this is essential for Pumilio-mediated protection of Xbp1 mRNA. These results identify an important regulatory mechanism connecting RBPs and the UPR.

## RESULTS

### The 3’UTR of Xbp1 contains cis elements that regulate mRNA stability

The 3’UTRs of *Drosophila* Xbp1^unspliced^ and Xbp1^spliced^ differ in the length, the former being 600bps longer than the latter (Fig. 1a). We conducted a search on the 3’UTR of *Drosophila* Xbp1 mRNA and identified two putative Pumilio regulatory elements (UGUAXAUA) (Fig. 1a,b). To test experimentally if these regions can regulate mRNA, we constructed reporters containing GFP, expressed under the control of the metalotheionin promoter^46,47^, together with the 3’UTR of Xbp1 in wild type or a mutated version of each of the two PREs (Fig. 1b). To examine the impact of these PREs on mRNA stability, we mutated the sequence encoding the PRE (UGUAXAUA) to the non-functional element ACAAXAUA (Fig. 1b).

**Figure 1.**
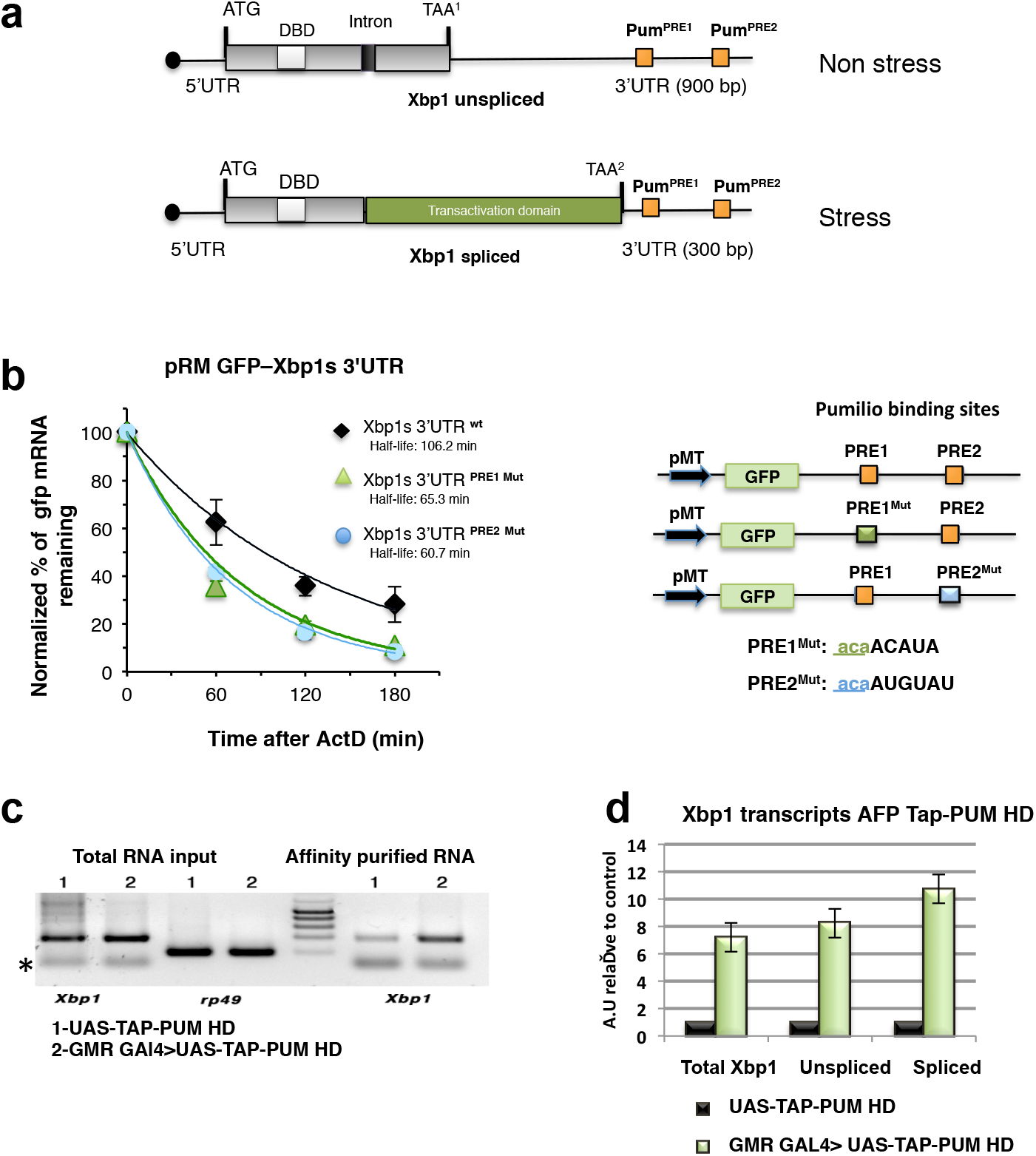
The 3’UTR of Xbp1 contains cis elements of mRNA stability. **a)** Schematic representation of *Drosophila* Xbp1 transcripts (unspliced and spliced). ATG (start codon), TAA (stop codon), DBD (DNA binding domain), Intron (23-nt hairpin structure recognized by IRE1 RNase domain. Two Pumilio regulatory elements (PRE1 and PRE2) are present in the 3’ UTR. **b)** Xbp1 3’UTR pRM GFP reporters bearing the WT and mutant PREs from the Xbp1 3’UTR. The consensus binding sites UGUA for Pumilio were mutagenized into ACAA for PRE1^mut^ and PRE2^mut^; pMT, metallothionein promoter. Reporters were transiently transfected in S2 cells. Total RNA was isolated at different points following the addition of actinomycin D (5 μg/ml). Stability of the GFP reporters was assessed by quantitative RT-PCR (qRT-PCR), using primers specific for GFP and rp49 mRNAs (control). Levels of mRNA reporter were normalized to those of rp49 mRNA, and averages and standard deviations from three independent experiments are plotted. **c)** TAP tag RNA affinity purification (AFP) from adult heads of transgenic flies expressing the Pumilio RNA-binding motif (GMR GAL4>UAS-TAP-PumHD) under the GMR driver (expression in eyes). Xbp1 transcripts are enriched in Pumilio RNA affinity purifications compared with controls (UAS TAP-PumHD, but no GMR GAL4). **d)** Quantitative RT-PCR using primers specific for Xbp1^Spliced^, Xbp1^Unspliced^ and total Xbp1 transcripts, from GMR GAL4>UAS-TAP-PumHD RNA pull downs, normalized against levels in the UAS-TAP-PumHD control pull downs. Results show a specific interaction between Xbp1^Spliced^ and Xbp1^Unspliced^ transcripts and PumHD.

The mRNA decay of the resulting reporters was analyzed in transiently transfected *Drosophila* S2 cells, which were treated with actinomycin D to stop transcription. For each mutated version of the PREs the stability of the GFP reporter decreased as compared to the wild-type 3’UTR, suggesting a role for these regulatory regions in the post-transcriptional regulation of Xbp1. We also compared the effect of the 3’UTRs of Xbp1^spliced^ and Xbp1^unspliced^ forms by quantitative RT-PCR, which showed a greater effect of the former 3’UTR on stability of the GFP reporter mRNA (Fig. S1). This result suggests that other, yet unidentified regulatory regions of RNA stability may be present in the 3’UTR of Xbp1^unspliced^.

### Pumilio binds Xbp1 mRNA in the developing *Drosophila* eye

To analyze if Pumilio could act as a regulator of Xbp1 in fly tissues, we expressed a transgene encoding a TAP-tagged^48^ Pumilio RNA-binding motif (UAS-TAP-PumHD), under control of the eye-specific GMR-GAL4 driver. We conducted TAP pull-down RNA affinity purifications (RPAs) from heads of transgenic adult flies with the genotype GMR– GAL4>UAS-TAP-PumHD, and performed RT-PCR to determine whether the endogenous Xbp1 mRNA was enriched in flies over-expressing TAP-Pum-HD. Xbp1 transcripts were strongly enriched in GMR-GAL4>UAS-TAP-PumHD flies as compared to UAS-TAP-PumHD controls (Fig. 1c,d). We found that Pumilio could bind equally well to the 3’UTR of Xbp1^unspliced^ and Xbp1^spliced^ (Fig. 1d). These results suggest that endogenous Xbp1 transcripts in the Drosophila eye are subject to regulation by Pumilio.

### Pumilio regulates the stability of Xbp1^spliced^ mRNA

The activity of Ire1 in mediating Xbp1 splicing can be assayed with an Xbp1-GFP reporter^49^, wherein Xbp1^spliced^ is tagged with GFP. We constructed a new Xbp1 reporter (Xbp1-HA-GFP) (Fig. 2a), which distinguishes the translation product of Xbp1^unspliced^, fused to an HA tag, from that of Xbp1^spliced^, fused to GFP. We first tested the expression of the reporter translation products by western blot under non stress and stress conditions in the presence of dithiothreitol (DTT), an ER stress inducing chemical (Fig. 2b). In this context, PUM overexpression led to an increase in Xbp1^spliced^, but not Xbp1^unspliced^ protein levels.

**Figure 2.**
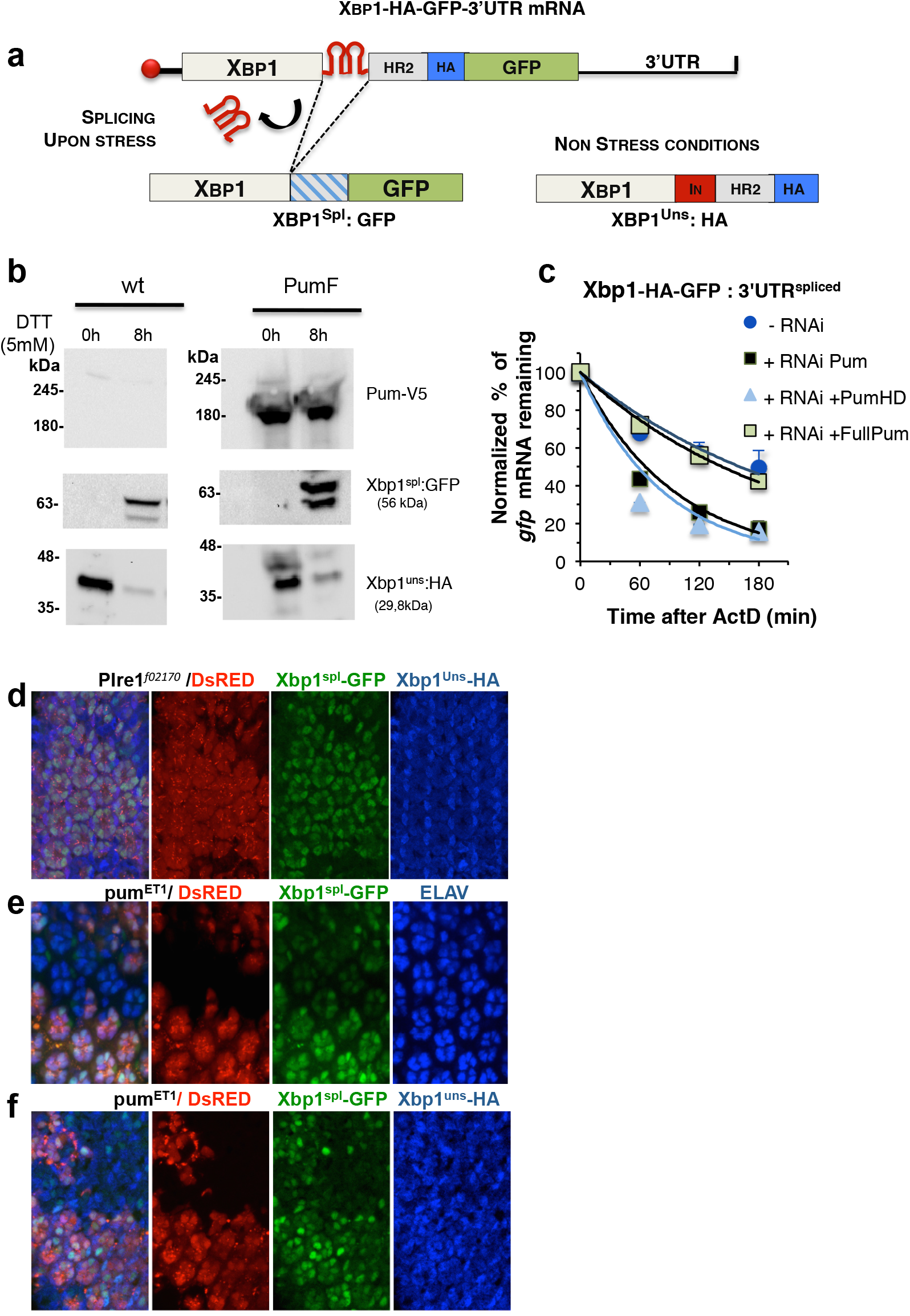
Pumilio regulates the stability of Xbp1^spliced^ mRNA. **a)** Schematic representation of the Xbp1-HA-GFP reporter construct. This reporter allows the detection of XBP1^Spl (Spliced)^ and Xbp1^Uns(Unspliced)^ forms. The pUAST-Xbp1-HA-GFP reporter, contains an HA tag fused in frame with the coding sequence of Xbp1^Unspliced^. GFP only gets in frame with Xbp1^Spliced^ after Ire1 mediated splicing of the 23bp intron. **b)** Expression of Xbp1-HA-GFP reporter in S2 cells under non stress (0h) and stress conditions (8h DTT treatment). Xbp1^spl^ GFP (56kDa) is detected under stress conditions. Overexpression of Pumilio (Pum-V5) increases the levels of Xbp1^spl^-GFP, but not of Xbp1^uns^-HA (29,8 kDa). **c)** Stability of the reporter was compared between non-treated S2 cells (-RNAi) and RNAi treated cells (+RNAi) against Pumilio. Inactivation of Pumilio by RNAi destabilizes Xbp1^spl^-GFP reporter (black squares). Reposition of Pumilio levels by transfection with full length Pumilio (Full Pum, green squares) restores the stability of the GFP reporter in S2 cells, in contrast to the RNA binding domain of Pumilio (PumHD, blue triangules). Total RNA was extracted after induction of stress with DTT (4h at 5μM) and mRNA stability was analysed by quantitative RT-PCR at defined periods after actinomycinD treatment (5μg/ml). **d)** The Xbp1-HA-GFP signaling reporter is activated in the *Drosophila* photoreceptors during midpupal stages (50 hrs). Xbp1^spl^-GFP is observed in WT photoreceptors (Red) but not in photoreceptors that are homozygous for an Ire1 null mutation (P*Ire1^f02170^*), labeled by the absence of DsRed. Xbp1^uns^-HA (blue) is observed in all cells. **e)** The expression of Xbp1^spl^-GFP (green) is reduced in cells homozygous for the Pum null mutation (PumET1), labeled by the absence of DsRed expression, in comparison with wildtype cells (DsRed positive). Elav (blue) was used as a marker of the photoreceptors. **f)** The expression Xbp1-GFP (green) is reduced but Xbp1 - HA (blue) is unaltered in cells homozygous for PumET1, labeled by the absence of DsRed expression, in comparison with wildtype cells (DsRed positive), indicating that the regulatory role of Pumilio is only on the Xbp1^Spl^ form. Anti-HA was used to label Xbp1^Uns^-HA protein.

Subsequently, we determined the stability of the Xbp1-HA-GFP reporter mRNA in transiently transfected S2 cells, either with or without Pumilio RNAi depletion (Fig. 2c). Depletion of Pumilio decreased the stability of the Xbp1-HA-GFP reporter transcript as compared with controls (Fig. 2c). When Pumilio levels were restored by co-transfecting S2 cells depleted of Pumilio with a plasmid expressing either the full-length Pumilio protein^50^ or the Pumilio RNA binding domain (Pum-HD), the mRNA stability of Xbp1-HA-GFP reporter returned to the levels of the control experiment (no RNAi treatment), but it did so only when the full-length protein was present (Fig. 2c). These results indicate that Pumilio has a protective role against degradation of the Xbp1 mRNA, and that although the HD region confers specific RNA interaction with the Xbp1 transcripts, the full-length Pumilio protein is required for its protective role.

### Pumilio regulates Xbp1^spliced^ during photoreceptor differentiation

Next we tested the regulation of Xbp1 mRNA by Pumilio in a physiological, developmental context. The Ire1 signaling pathway is activated during the pupal stages in the photoreceptors^51^, where it is required for photoreceptor differentiation and morphogenesis of the rhabdomere – the light-sensing organelle of photoreceptors. We generated transgenic flies with the Xbp1-HA-GFP reporter, and performed immunofluorescence analysis with antibodies against GFP and HA, to assess the expression levels of Xbp1^spliced^-GFP and Xbp1^unspliced^-HA in eyes containing clones for an Ire1 null mutation^51^. As expected, and validating the Xbp1-HA-GFP reporter, Xbp1^spliced^-GFP expression was absent from cells homozygous for Ire1 mutation (labeled by the absence of DsRed), but Xbp1^unspliced^-HA expression was unaffected in Ire1 mutant cells (Fig. 2d). We next examined the expression of Xbp1-HA-GPF in eyes containing clones of the Pumilio null mutation pumET1^52^. In pupal eyes (50 hr), in the absence of Pumilio (labeled by the absence of DsRed), the expression of Xbp1^spliced^-GFP was reduced as compared to the wild type cells (Fig. 2e,f). However, expression of Xbp1^unspliced^-HA remained unchanged (Fig. 2f), indicating that Pumilio exerts its regulatory role on the mRNA of Xbp1^spliced^-GFP but not Xbp1^unspliced^-HA

### Pumilio undergoes IRE1 kinase-dependent phosphorylation during ER stress

It is known that Pumilio proteins may change their activity according to their phosphorylation state^53^. Furthermore, the observation that the stability of the Xbp1^spliced^ transcript was dependent of the full-length Pumilio protein (Fig. 2c) indicated that other domains of Pumilio, besides the RNA-binding domain, are also important for Xbp1 mRNA stability. Recent studies^36,42,50^ indicate that additional regions in the N terminus of Pumilio act as repressor domains, with two specific segments – designated Pumilio conserved motif (PCM) a and b – being found in both Drosophila and human PUMs (Fig. 3a and Fig. S2a). We screened the Pumilio protein sequence for potential serine/threonine phosphorylation sites using online bioinformatic tools^54^ combined with phosphorylation sites reported in other screens (http://www.phosphosite.org/). We found potential phosphorylation sites across the different domains of Pumilio (Fig. 3a). To investigate if Pumilio is phosphorylated during ER stress, we conducted Phostag analysis using Pumilio proteins tagged with a C-terminal V5 epitope (Fig. 3a). It was difficult to analyze phosphorylation of full-length Pumilio^50^, due to its relatively large size of ~180 kDa (data not shown). Therefore, to facilitate the analysis, we constructed 3 truncated versions of V5-tagged Pumilio (D1= aa 1 to 547; D3= aa 77 to 1091 and D1D3= aa 1 to 1091), which contained the predicted phosphorylation sites.

**Figure 3.**
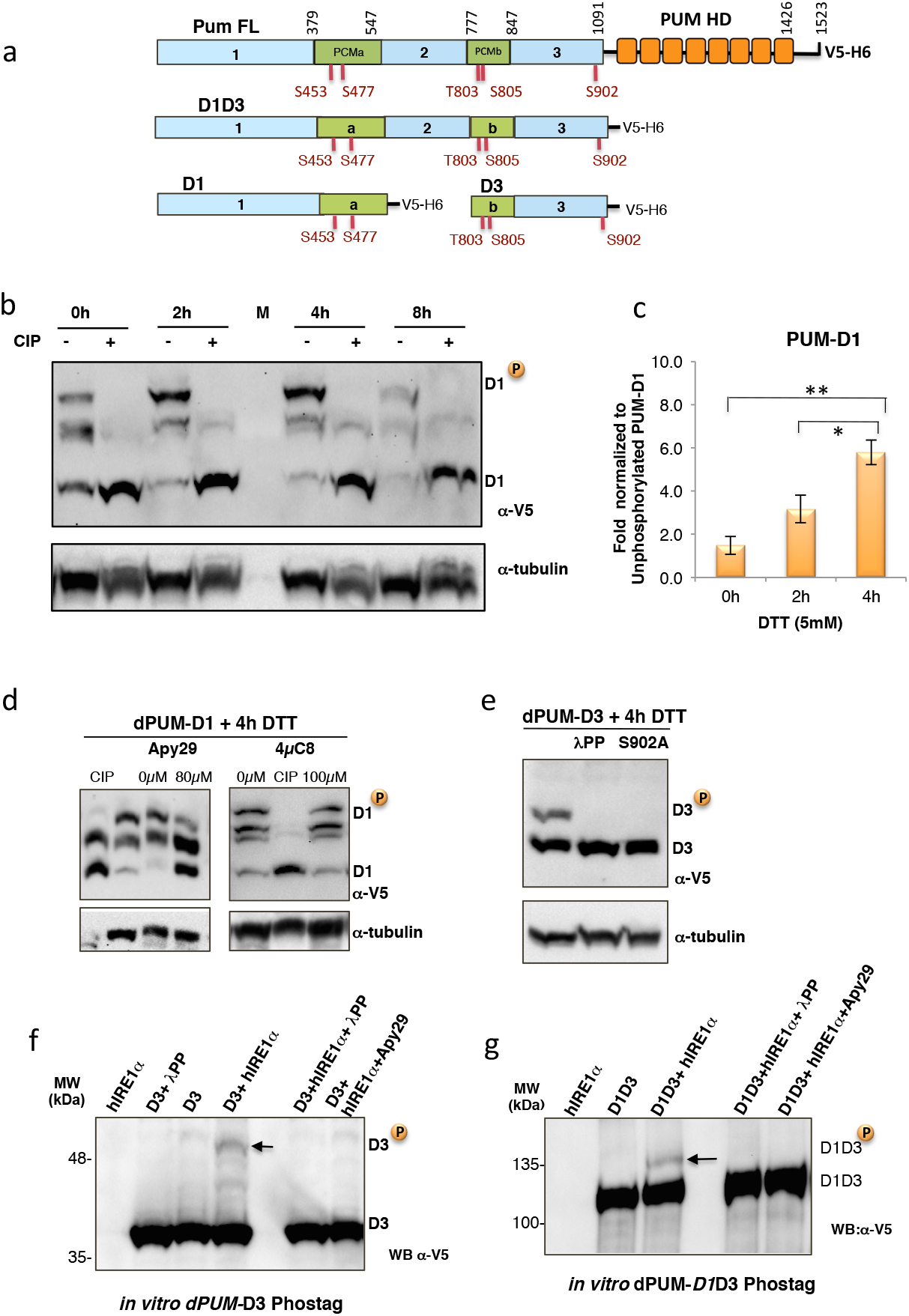
Pumilio undergoes Ire1 kinase-dependent phosphorylation during ER stress. **a)** Diagram of *Drosophila* Pumilio domains, with possible phosphorylation sites identified in Phosphosite (www.phosphosite.org and Phosida (http://www.phosida.com). Indicate below in red for each amino acid residue. Truncated version of the full-length Pumilio protein were constructed carrying different domains of Pumilio (Pum FL: Full length protein, D1D3: domain 1 to 3, D1: domain 1, D3: domain 3, PUM-HD: RNA binding Pumilio homology domain. Constructs are tagged with V5 and 6xHis at their C-terminal end. The motifs represented by letters a and b refer to conserved motifs (PCMa and PCMb) in *Drosophila* and human PUM proteins. **b)** *Drosophila* S2 stable cell lines expressing the V5 tagged Pumilio D1 were submitted to ER stress (DTT 5mM) and samples collected at defined times before (0h) and after stress induction (2h, 4h 8h). Phosphorylation of Pumilio domains was determined based on changes in their electrophoretic mobility through a standard SDS-PAGE gel using the Phostag compound [50mM]. Pumilio proteins were detected by immunoblot (IB) with a mouse monoclonal V5 antibody. The mobility shift of Pumilio is specific for phosphorylation as it is reversed by calf intestinal phosphatase treatment (+CIP). Total protein levels were monitored by tubulin levels. D1^P^: hyper-phosphorylated forms, D1: non phosphorylated form of D1. M: protein size markers. **c)** Quantification of fold increase of PUM D1^P^ during ER stress, normalized to non phosphorylated form (D1). The statistical significance was calculated by one-way ANOVA coupled with Tukey’s post hoc test (* p<0.05; ** p<0,01) **d)** An inhibitor of Ire1 kinase activity impairs the phosphorylation of dPUM-D1. Immunoblot (IB) with Phostag phosphorylation analysis of Pumilio D1 after ER stress (4h DTT treatment). Cells were incubated with an inhibitor of Ire1 kinase activity (APY29, 80μM) or an inhibitor of IRE1 RNase activity (4μ8c, 100μM). The phosphorylation pattern of dPUM-D1 was monitored by IB with anti-V5 antibody. Total protein loading was monitored by levels of tubulin. CIP (alkaline phosphatase treatment of cell extracts after 4h of DTT treatment. **e)** Phostag immunoblot (IB) for phosphorylation analysis of Pumilio Domain 3 (dPUM-D3) after ER stress (4h DTT treatment). dPUM-D3 was detected by IB using anti-V5 antibody. Total protein loading was monitored by levels of Tubulin. Mutation of site Ser902 to Ala (S902A) prevents the DTT induced phosphorylation of dPUM-D3. λPP: λ phosphatase treatment of cell extracts after 4h of DTT treatment. **f, g)** Phostag immunoblot (IB) after in vitro phosphorylation assay with purified domains of Pumilio (D3 and D1D3) incubated with human IRE1-alpha [464-977] in kinase buffer (2mM ATP). Controls: IRE1 alone, PUM alone (D3 or D1D3); Inhibition of phosphorylation was observed upon treatment with λ-phosphatase or the Apy 29 Ire1 kinase inhibitor. Pumilio proteins were detected with anti-mouse V5 antibody.

Phostag analysis revealed that, during ER stress (4 hr DTT), Pumilio D1 displayed a 6-fold increase in phosphorylated forms, relative to non-ER stress conditions (0 hr DTT) (Fig. 3b). This phosphorylation pattern was confirmed by treatment with phosphatases (+CIP or + λPP), which reverted Pumilio to a non-phosphorylated state (Fig. 3b).

During ER stress and after Ire1 trans-autophosphorylation, the cytosolic kinase-endoribonuclease domain of IRE1 rotates to a back-to-back configuration^2,18^, possibly allowing the kinase active site to access other potential substrates. We reasoned that through binding to the 3’UTR of Xbp1, Pumilio may localize to the vicinity of the Ire1 kinase domain. Therefore, we tested whether Ire1 could phosphorylate Pumilio during ER stress, using the inhibitors of Ire1 kinase activity (Apy 29)^55^ and compound #18^56^ or an inhibitor of IRE1 RNase activity (4μ8C)^57^. Upon DTT exposure (4 hr), in S2 cells transfected with PUM-D1, Apy 29 attenuated the generation of PUM-D1 hyper-phosphorylated forms (Fig. 3d). In contrast, 4μ8C did not change the pattern of PUM-D1 phosphorylation, which is expected as 4μ8C is specific for the RNase activity and does not inhibit IRE1 kinase activity^57^. We also observed ER stress induced phosphorylation of PUM-D3, and by constructing phosphorylation site mutant versions of PUM-D3 we could identify S902 as a site that is phosphorylated under ER stress conditions (Fig. 3e).

### hIRE1 phosphorylates Pumilio and hPUM1

To confirm that IRE1 kinase activity could directly phosphorylate Pumilio, we performed *in vitro* kinase reactions using the hIRE1alpha kinase-endoribonuclease (KR) domain^57^ and purified versions of Pumilio domains D3 and D1D3. Phostag immunoblot analysis (Fig. 3f,g) showed specific bands indicating phosphorylation of PUM-D3 and PUM-D1D3 upon incubation with hIRE1 KR. Pumilio phosphorylation was abolished upon Apy 29 or λ-PP treatment. To complement these results, we performed *in vitro* radioactive kinase assays using hIRE1 KR and purified versions of Pumilio and hPum1 protein domains (Fig. 4a,b and Fig. S2b). The hIRE1 KR protein phosphorylated both Pumilio and hPum1 proteins. For negative controls, we used a kinase-dead hIRE1 KR as enzyme, or BIP (a luminal ER protein that should not be a direct target for IRE1 phosphorylation) as substrate (Fig. S3a). Furthermore, to assure that the phosphorylation pattern observed for the PUM proteins in the radioactive assays did not simply reflect IRE1 auto-phosphorylation, we used an antibody specific for phosphorylated IRE1. While we observed phosphorylated monomers, dimers and oligomers of hIRE1 KR as expected, we could not detect any signal in the 40 KDa region (Fig. S3b), which corresponds to the phosphorylated forms of Pumilio that were detected with the V5 antibody (Fig. 4b). These results show that hIRE1 KR can directly phosphorylate Pumilio.

**Figure 4.**
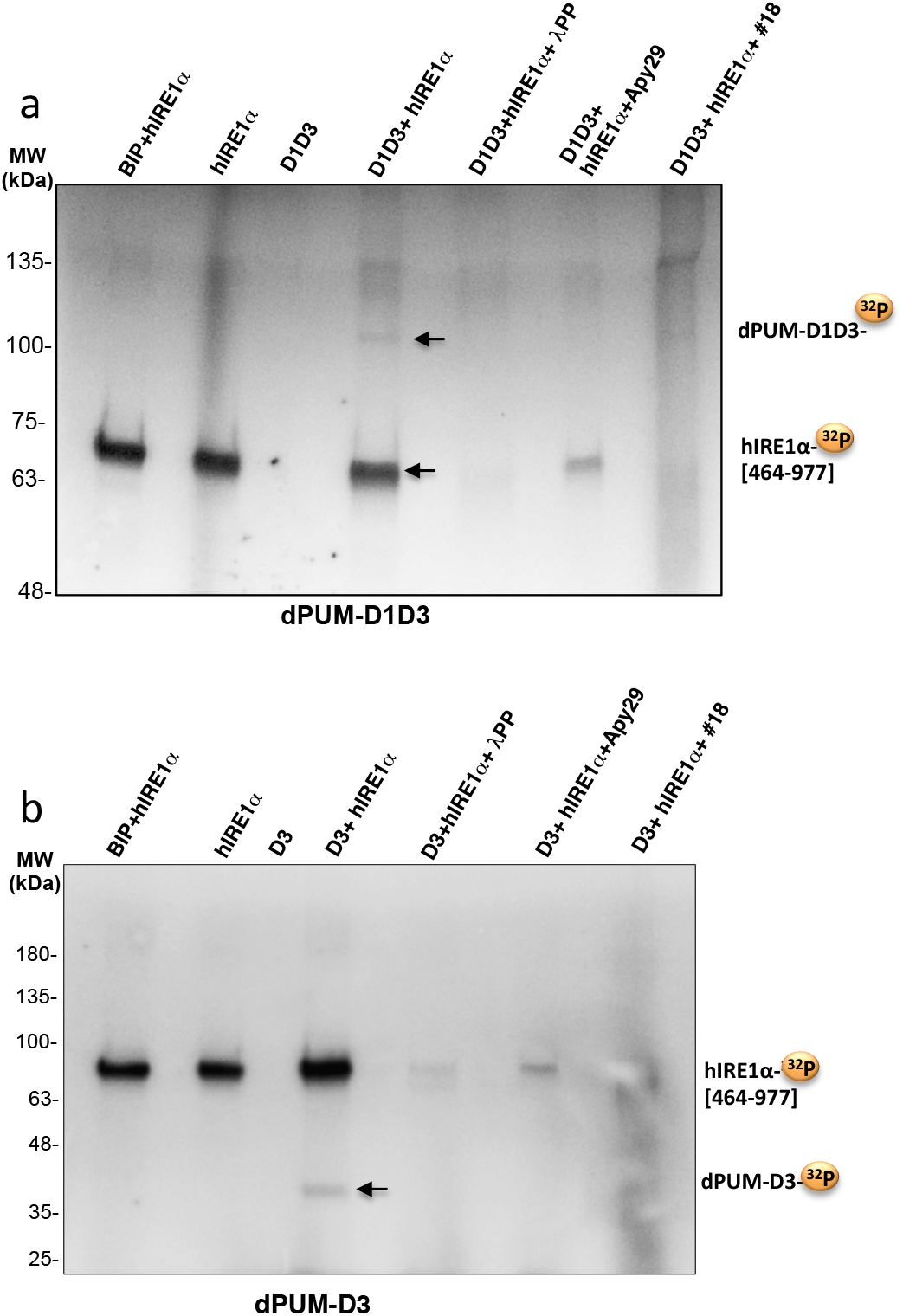
hIRE1 KR phosphorylates Pumilio in vitro. **a** and **b**) *In vitro* radioactive phosphorylation assays of purified Pumilio dPUM-D1D3 (a) and dPUM-D3 (b) with hIRE1 KR (464-977). Purified proteins were incubated 2h in IRE1 kinase buffer containing γ-ATP[^32^P]. Detection of radioactive dPUM-D1D3, dPUM-D3 and auto-phosphorylation of hIRE1 KR are denoted by a yellow bullet ^32^P. Negative controls: BIP as a luminal ER protein non-phosphorylated by IRE1, and proteins without incubation with IRE1. Specificity of phosphorylation was monitored by treatment with lambdaPP or incubation with specific inhibitors of IRE1 kinase activity (Apy 29 and compound #18). MW: protein molecular weights in kDa.

### Pumilio protects Xbp1 mRNA from regulated IRE1-dependent decay

We have shown that Pumilio has a protective role in the stability of Xbp1 mRNA (Fig. 2c). Ire1 can also mediate RIDD, the selective decay of ER-bound mRNAs, thereby reducing the load of nascent proteins entering the ER in order to be folded^58^. RIDD plays an import role during persistent ER stress^19^ and in *Drosophila* cells it can lead to the complete degradation of even ectopic mRNAs, such as GFP^15,59^. We therefore hypothesized that Pumilio may protect Xbp1 mRNA from decay by Ire1 under ER stress conditions. To test this possibility, we performed mRNA stability assays with the Xbp1-HA-GFP reporter in S2 cells treated with DTT, in the absence or presence of the Ire1 RNase inhibitor 4μ8C. The destabilization of Xbp1-HA-GFP mRNA in cells depleted of Pumilio was reverted upon 4μ8C treatment (Fig. 5a), indicating that Pumilio stabilizes Xbp1 mRNA against the RNase activity of Ire1.

**Figure 5.**
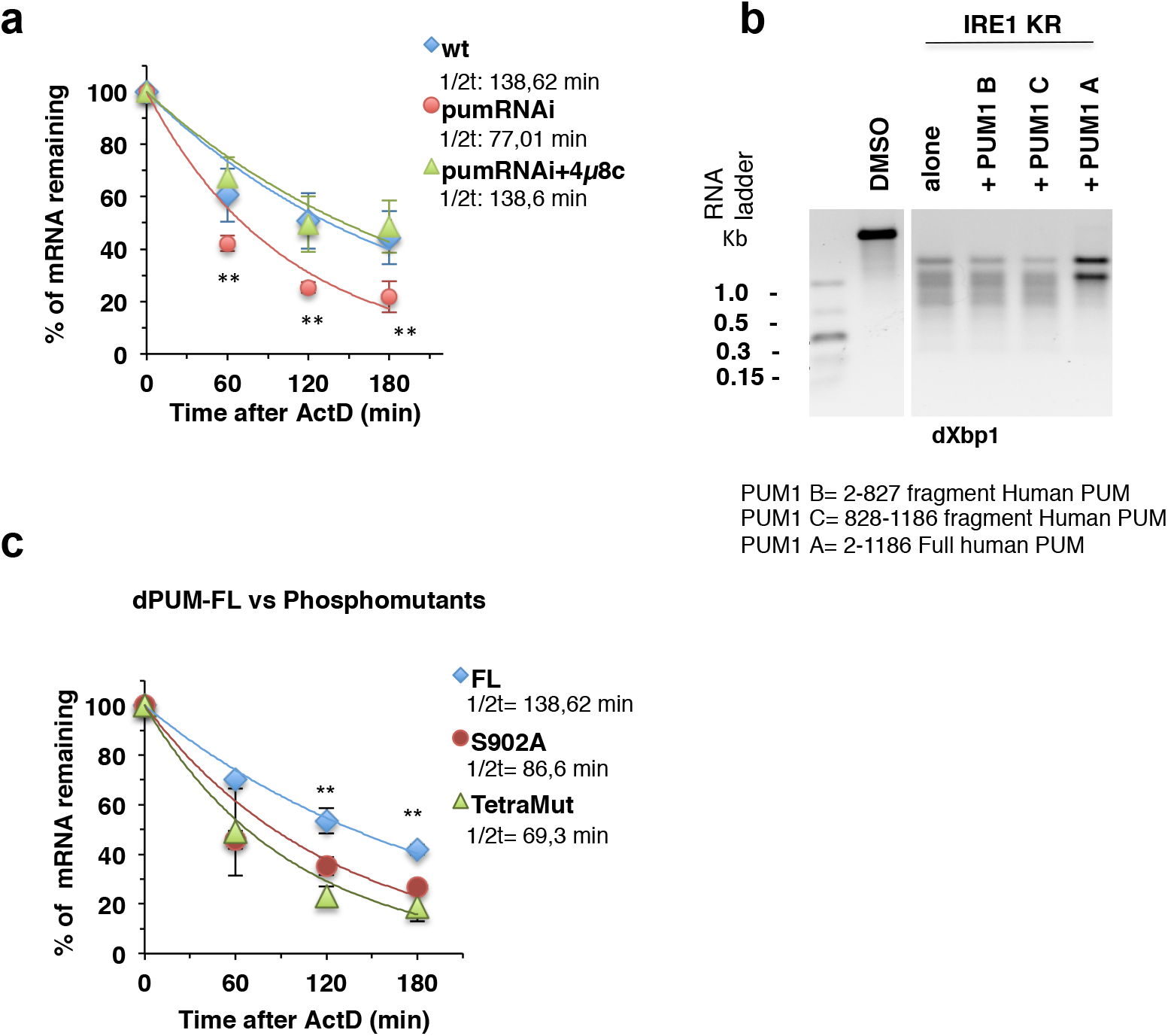
Pumilio protects Xbp1^spliced^ mRNA from regulated Ire1-dependent decay. **a)** WT S2 cells were compared to cells treated with RNAi against Pumilio (pumRNAi) and with or without 4μ8c [100 μM], an inhibitor of IRE1 RNase activity. Following ER stress induction (5mM DTT), and in the presence ActinomycinD [5μg/ml], total RNA was isolated from cell extracts at different periods. The levels Xbp1^spliced^-GFP mRNA were normalized to *rp49* mRNA levels, and the half-lives calculated by regression analysis of amount of %mRNA remaining against time (min). qRT-PCR analysis was used to determine mRNA levels using specific primers for *gfp* and *rp49*. Averages of % mRNA remaining and corresponding standard deviations were calculated from at least 3 independent experiments. The statistical significance was calculated by one-way ANOVA coupled with Tukey’s post hoc test (* p<0.05; ** p<0,01). **b)** *In vitro* transcripts of dXbp1 were incubated with purified hIRE1 KR. Human PUM1 FL protects Xbp1 RNA from Ire1 dependent decay, but does not impair Ire1 dependent Xbp1 splicing. dXBP1 = *Drosophila* XBP1 T7 transcript, hPUM1 B = [2-827] fragment Human PUM; hPUM1 C=[828-1186] fragment Human PUM, hPUM1 A= [2-1186]; 3P-IRE1 = phosphorylated IRE1. **c)** qRT-PCR results from Pumilio RNai treated S2 cells transfected with plasmid expressing dPUM-FL, a single or a quadruple phosphomutant of dPUM-FL (full length Pumilio containing the single mutation S902A; and a quadruple mutant: T537A, S540A, S544A, S902A). The Pumilio protective role upon Xbp1^spliced^-GFP is diminished in the dPUM-FL phosphomutants. The statistical significance was calculated by one-way ANOVA coupled with Tukey’s post hoc test (* p<0.05; ** p<0,01).

We also conducted mRNA stability assays using *in vitro* transcribed *Drosophila* Xbp1 mRNA incubated with hIRE1 KR. Since we could not produce purified recombinant full-length Pumilio, we used different forms of hPUM1, which were pre-incubated with the Xbp1 transcript in order to assess their protection against degradation by IRE1 (Fig. 5b). Full-length hPUM (hPUM1-FL) protected the Xbp1 mRNA from degradation by hIRE1 KR, but did not block or even diminish the hairpin motif-dependent splicing of Xbp1 mRNA. By contrast, hPUM1-B (2-827) did not protect Xbp1 mRNA from decay, consistent with the absence of the HD region necessary for binding to Xbp1 mRNA in this hPUM variant. Likewise hPUM1-C (828-1186) is incapable of such protection. Only the full-length hPUM1 protein contains all the requisite domains for protection of Xbp1 mRNA, in keeping with the above results (Fig. 2c).

### Pumilio’s protection of Xbp1 mRNA is dependent on its phosphorylation state

Having identified S902 as a site of Pumilio phosphorylation under ER stress (Fig. 3e), we next asked if the phosphoryation status of Pumilio could regulate the effect on the stability of Xbp1^spliced^. To examine this, we compared the half-life of Xbp1 upon overexpression of wt Pumilio or Pumilio phosphomutants (S902A and a quadruple mutant: S902A,T537A, S540A, S544A) in RNAi Pumilio-depleted S2 cells (Fig. 5d). As compared to wild type Pumilio, overexpression of the phosphomutants lead to a decrease in the stability of Xbp1^spliced^, demonstrating that Pumilio phosphorylation is required to promote its protective effect on Xbp1 transcripts under ER stress.

## DISCUSSION

Our results uncover a novel protective effect of Pumilio on Xbp1 mRNA during ER stress. This protective effect depends on the phosphorylation status of Pumilio, which can be mediated by Ire1 kinase activity in response to ER stress. We propose a model depicted in Fig. 6, which involves 3 steps.

**Figure 6.**
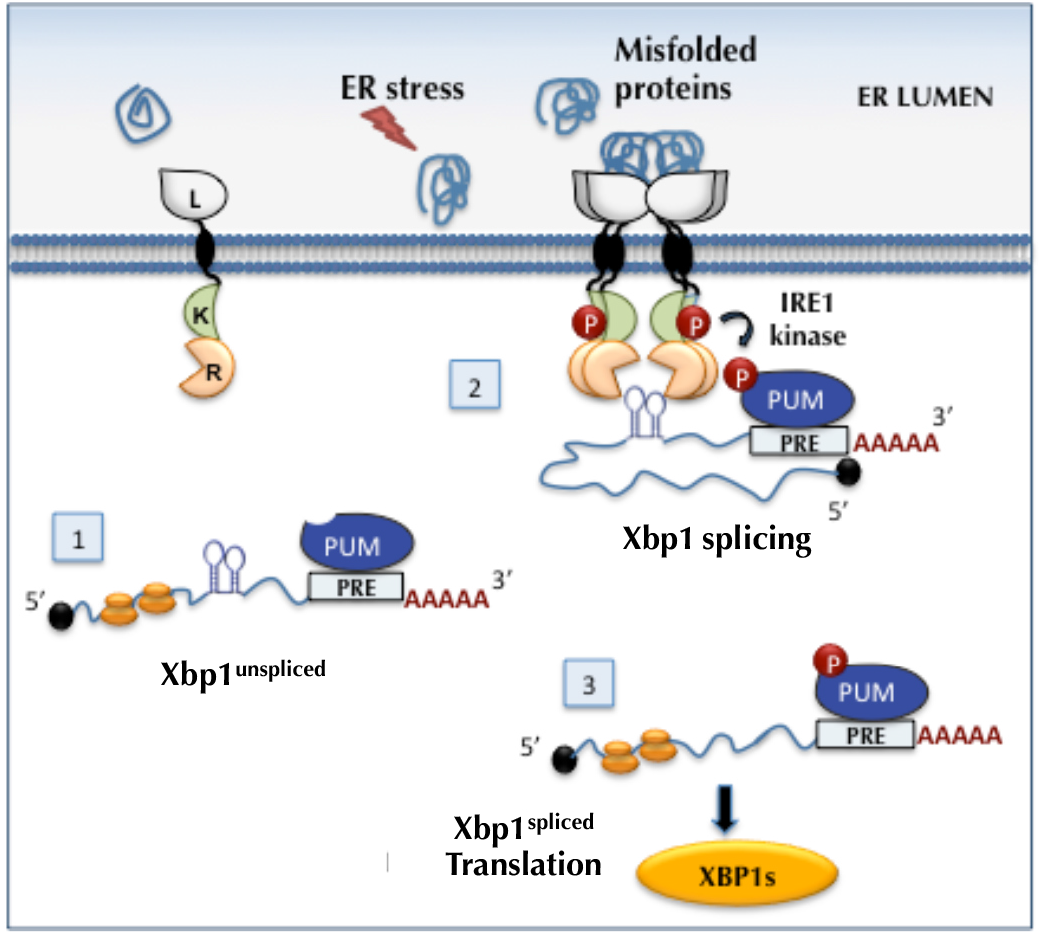
Model for Ire1-controlled post-transcriptional regulation of Xbp1 by Pumilio. Step 1-In non ER stress conditions, Xbp1^unspliced^ is present in the cytosol and PUM may bind to the PRE elements in the Xbp1 3’UTR; Step 2-ER stress conditions lead to Ire1 autophosphorylation and Pumilio phosphorylation. Phosphorylated Pumilio protects Xbp1^spliced^ mRNA from Ire1 dependent decay, but does not impair Ire1 dependent Xbp1 splicing. Step 3 - Xbp1^spliced^ transcripts can be efficiently translated into the Xbp1 ^spliced^ transcription factor.

In the absence of ER stress, Ire1 is not active and Xbp1 mRNA is not spliced, but Pumilio can bind the 3’UTR of Xbp1^unspliced^ mRNA (Fig. 6 - 1). Indeed, our RNA pull down experiment (Fig. 1d) showed that Pumilio can bind equally well to the 3’UTR of Xbp1^unspliced^ and Xbp1^spliced^. Accumulation of misfolded protein in the lumen of the ER causes ER stress and activates Ire1. At this stage, in order to be spliced, Xbp1 mRNA must localize in the vicinity of the cytoplasmic domain of Ire1, which harbors both kinase and endoribonuclease activity. Upon ER stress, Pumilio is phosphorylated via a mechanism that requires Ire1 kinase activity (Fig. 3). As demonstrated by our *in vitro* assays (Fig. 4), the kinase domain of Ire1 can directly phosphorylate Pumilio; however, we cannot exclude at this stage the possibility that other kinases also may contribute to Pumilio phosphorylation. It is known that phosphorylation of Pum family proteins contributes to their activation and function^53^. We suggest that in this case, Ire1-dependent phosphorylation of Pumilio causes a conformational change in the mode of interaction between Pumilio and Xbp1 mRNA, such that much of the transcript, though not the canonical stem-loop structures, is inaccessible to the more promiscuous endoribonuclease activity of Ire1 (Fig. 6: step 2). This allows Xbp1 splicing to occur and furthermore permits Ire1 to carry out RIDD while sparing Xbp1^spliced^ mRNA and allowing its more efficient translation (Fig. 6 - 3).

We do not know if Pumilio also protects Xbp1 mRNA from other ribonucleases and for how long the protective effect against RIDD lasts. Other studies have shown^60^, using cultured mouse cells, an increase in the stability of mRNA Xbp1^spliced^ at a time of translational repression during “early” phases of ER stress (4 hr treatment with DTT). These authors suggested that Xbp1^spliced^ mRNA could be protected from degradation by an unidentified protein factor. Our results suggest that Pumilio is such factor, while Ire1 RIDD activity is the degradation-causing agent.

From our model, we also predict that upon overcoming ER stress conditions, Pumilio phosphorylation levels will be reduced, either due to the activity of phosphatases and/or due to Ire1 dephosphorylation and attenuation^61^. At this stage, Pumilio’s protective role over Xbp1 mRNA may subside, presumably making the transcript more vulnerable to the action of the mRNA decay machinery and the ribosome-associated quality control, as previously shown for Hac1^62^. This regulatory step may avoid an accumulation of Xbp1^spliced^ under non stress conditions, which could be detrimental for cellular homeostasis.

Previous studies have shown that Pumilio proteins play a translational repressive role or promote degradation of their target mRNAs, which is in contrast with our results. However, another study has shown that, contrary to the canonical repressive activity, PUM1/2 rather promote FOXP1 expression through direct binding to two consensus PREs present in the FOXP1-3’UTR^63^. Furthermore, an additional report has identified RNAs that are positively regulated by human PUM1 and PUM2^64^.

Our results demonstrate that Ire1 is responsible for the phosphorylation status of Pumilio during ER stress and that this phosphorylation has implications for Xbp1^spliced^ protein levels. Recent data in zebrafish implicate Pum1 phosphorylation as an initial key step for the sequential activation of cyclinB1 mRNA translation during oocyte maturation, although the kinase involved remains unidentified ^65^. In Xenopus, Nemo-like kinase (NLK) – a typical mitogen-activated protein kinase that is activated during an early phase of oocyte maturation^66^ – was shown to directly phosphorylate Pum1 as well Pum2 *in vitro*. It is possible that other kinases may be involved in Pumilio phosphorylation under non-ER stress conditions or specific developmental stages.

In conclusion, our present work uncovers an unanticipated mechanism that regulates one of the key branches of the UPR through the action of an RBP. This involves Ire1-kinase-driven phosphorylation of Pumilio, which in turn protects Ire1-RNase-driven Xbp1^spliced^ against RIDD, thereby coordinating the two major endoribonuclease outputs of IRE1 to enable a more efficient intracellular response and ER stress mitigation.

## METHODS

### Cell culture

*Drosophila* S2 cells (Schneider, 1972) were cultured at 25 °C in Schneider’s *Drosophila* medium (Invitrogen) supplemented with 10% heat inactivated fetal bovine serum (FBS, Invitrogen), mM glutamine (Invitrogen), 100 U/mL penicillin and 100 μg/mL streptomycin (Invitrogen).

### Transgenic flies

Transgenic lines were generated using pUAST-AttB and phiC31 integrase-mediated DNA integration (BestGene) that allows the insertion of the transgenes in a specific site of the acceptor fly genome. Clones of mutant eye tissue were generated by the Flp/FRT technique^67^, with Flipase expression under the control of the *eyeless* promoter. Drosophila stocks obtained from the Bloomington Stock Center (Indiana University, Bloomington, IN, USA): GMR-Gal4 (active in the eye, under the control of the glass multiple reporter); eye-flip GMR-Gal4 (promotes recombination in the eye) and Actin5c-Gal4 (ubiquitous expression). *Drosophila* stocks were maintained at 25°C on standard cornmeal media in an incubator with a 12h light/dark cycle. TAP-PUM stocks were a kind gift from André Gerber Laboratory.

### Plasmid construction

The 3′UTRs of Xbp1^spliced^ and Xbp1^unspliced^ forms were amplified from the cDNA clone GH09250 (Flybase) using specific primer pairs, and cloned downstream of the green fluorescent protein (GFP) coding sequence in the vector pRmHa, containing the metallothionein promoter. The mutations in the Pumilio binding sites were introduced using oligonucleotide-mediated site-directed mutagenesis and inverse PCR. The UAS-Xbp1-HA-GFP-3’UTR constructs were made by PCR cloning. All cloning was performed with the Phusion High-Fidelity PCR Master Mix with HF Buffer or GH Buffer according to the manufacturer’s protocol. All clones were confirmed by sequencing (Stabvida). The full-length Pumilio protein with a C-terminal V5 tag^50^ was a gift from Aaron Goldstrohm, from which the different truncated dPUM-D1, dPUM-D3 and dPUM-D1D3 were constructed.

### Transfection and stable cell line establishment

Cells were co-transfected with the different plasmids using Effectene reagent according to manufacture indications (Qiagen). For stable transfection, cells were selected by replacing Schneider’s complete *Drosophila* media with fresh medium supplemented with the appropriate antibiotics (zeocin, puromycin) according to the resistance gene present in the transfected plasmids and maintained under selective media until the formation of resistant clones.

### Total RNA and protein extraction

For RNA stability and Western blot experiments, 4,5×10^5^ S2 cells were seeded in 12-well plates the day before treatments, transfections and protein or RNA extraction. Transfection was performed with Effectene (Qiagen) according to the manufacturer’s instructions. Expression of the *GFP* reporter under the metallothionein promoter was induced by adding 7 mM CuSO_4_ to the cell culture media for 3hrs. For the UAS dependent Xbp1 reporters, transcription was induced by co-transfection of Actin-Gal4 plasmid. For mRNA half-life measurements, transcription was blocked with actinomycin D (5 μg/ml; Sigma-Aldrich) and the cells were harvested at the indicated time points. Total RNA was then extracted using Zymo Research quick-RNA miniprep Kit.

### qPCR and RT-PCR

Primers for qRT–PCR analysis were designed according to MIQE guidelines (Minimum Information for Publication of qRT-PCR Experiments) using NCBI primer blast, choosing a melting temperature of 62°C. By using three serial dilutions of cDNA, primer efficiencies were determined, only primers with efficiencies varying around 100% were used for analysis. 0,5 μg of total RNA was retro-transcribed using RevertAid H Minus First Strand cDNA Synthesis Kit (Thermo/Fermentas). Each PCR reaction was performed on 1/40 of the cDNA obtained, using SSoFast EvaGreen Supermix (Bio-Rad) according to the manufacturer’s instructions and Bio-Rad CFX-96 as detection system. All samples were analyzed in triplicates and from 3 independent biological RNA samples. For each sample, the levels of mRNAs were normalized using rp49 as a loading control. Normalized data then were used to quantify the relative levels of mRNA using the ΔΔ*CT* method. qPCR was carried out using a CFX-96 Biorad instrument. Biological replicates represent independently grown and processed cells. Technical replicates represent multiple measurements of the same biological sample.

### Protein analysis

Total protein lysates were prepared in lysis buffer containing protease inhibitors. Proteins were size-separated by SDS-PAGE and transferred onto nitrocellulose or PDVF membranes (Biorad). For Western blot analysis, primary antibodies were anti-GFP (3H9) (1:1000, Chromotek), anti-mouse V5 (1:5000,Invitrogen), anti-mouse HA (1:5000, Covance) and anti-α-tubulin (AA4.3) (1:1000, Developmental Studies Hybridoma Bank).

### Protein Purifications

Drosophila Pumilio domains (PUM-D1, PUM-D3, Pum-D1D3) fused to V5-6xHis were subcloned into pET 28a and pET26a vectors for protein expression in bacteria. Plasmids were transformed into BL21(DE3) competent cells and recombinant proteins were induced with 1mM IPTG at 25°C. Bacteria were lysed in lysis buffer (500 mM NaCl, 50 mM Tris-Cl, pH8.0, 0,1% Triton, Protease inhibitors). Samples were lysed by sonication, centrifuged twice at 12,000rpm for 15min, and the cleared supernatant was bound to Ni-NTA Superflow beads (Qiagen) by gravity filtration. Unbound proteins were washed with lysis buffer at pH 8.0 supplemented with increasing amounts of imidazole (5mM, 40mM, 60mM). Recombinant proteins were eluted from beads in lysis buffer with 100-250 mM imidazole at pH 8.0. For recombinant proteins retained in inclusion bodies (PUM-D3), solubilization was done by including 6M urea in lysis buffer. All purified Pumilio recombinant proteins were dialized overnigth at 4°C, against a final buffer (20mM Hepes, 200 mM NaCl, 5% Glicerol, 2mM DTT) and concentrated with appropriate MW cut-off Vivaspin columns (Merck). The concentration of purified proteins was determined by colorimetric assay (Bio-Rad DC Protein Assay) and verified by electrophoresis alongside of BSA standards with Coomassie staining.

### Immunoprecipitation of TAP-PUM from flies

Extracts from adult flies heads were prepared as described in ^48^. After immunoprecipitation of TAP-PUM, total RNA was extract with Trizol and Xbp1 mRNA levels detected by RT-PCR using specific primers for each Xbp1 transcript.

### In vitro phosphorylation assays

Ire1 phosphorylation assays were performed by incubation of purified hIre1 KR (3,3μg/μl) with purified Pumilio protein (dPumD1, dPumD3, hPumFl) in IRE1 kinase buffer (25mM HEPES, pH 7.5, 150mM NaCl, 5mM DTT, 5% glicerol), containing either 10μCi of γ-ATP[^32^P] for radioactive assays or cold ATP [2mM] for phostag immunoblot assays (in a total volume of reaction of 20μl). Inhibitors of IRE1 Kinase (Apy 29 [2,5μM/μl] and compound #18 [2,5μM/μl]) were pre-incubated with hIRE1 and all reactions were assembled on ice, prior to addition of ATP and incubation for 2h at 25°C. Phosphatase treatment with λPP (NEB) or CIP (NEB) were performed after phosphorylation reactions, for 40 min at 30°C. Each reaction was stopped by addition of SDS-PAGE loading buffer and run on pre-maid SDS-PAGE gels (Biorad). The autophosphorylation of hIRE1 KR was confirmed by western blot using a pSer/24 phospho-specific antibody (Genentech) and phosphorylated Pumilio proteins were detected using anti-mouse-V5 antibody (Invitrogen). In the case of kinase radioactive assays, gels were dried and kinase activity visualized by autoradiography.

### Phostag-gels

S2 cells were lysed in CIP buffer (100 mM NaCl, 50 mM Tris-HCl pH 7.9, 10 mM MgCl2, 1 mM DTT), 1 mM PMSF, 0.1% NP40, protease inhibitor cocktail (Roche), and phosphatase inhibitor cocktail (Calbiochem). For CIP treatment phosphatase inhibitor cocktail was omitted, and lysate was incubated 37°C 60 min in 2 units CIP (NEB) per 50 μL reaction containing 50μg of total protein. For λ-phosphatase (NEB) treatment, lysates were incubated with 1 unit of phosphatase for 30 min at 30°C in λ phosphatase buffer. Lysates were cleared by centrifugation and subjected to SDS-PAGE. To detect phosphorylated Pumilio in SDS-PAGE, we used Phos-tag AAL-107 (Wako Chemicals GmbH) according to the manufacturer’s instruction ^68,69^. Western blotting was performed using mouse anti-V5 (1:5000, Invitrogen), followed by the corresponding Horseradish Peroxidase (HRP) conjugated secondary antibodies (1:5000, GE Healthcare) and visualized using the ECL Plus Western Blotting detection system (GE Healthcare).

### dsRNA treatments

RNAi was performed as described previously ^70^. Primer pairs tailed with the T7 RNA polymerase promoter were used to amplify PCR fragments obtained from cDNA clones. PCR products with an average size of 600 bp were then used as templates for dsRNA production with the T7 RiboMAX system (Promega). For transfection, 15 μg/ml dsRNA was added to S2 cells in 12-well plates, during 9 days to silence *Drosophila* Pumilio. mRNA depletion was confirmed by RT-PCR before further transfections.

### Immunofluorescence and confocal microscopy

For Drosophila pupal dissections, white pre-pupae (0h pupa) were collected and maintained at 25°C until the required stage. Larval, pupal and adult eyes were dissected in 1xPBS, fixed in 1xPBS + 4% Formaldehyde for 40 minutes at room temperature and washed 3 times with 1xPBS + 0.3% Triton X-100. Primary antibodies were incubated in 1xPBS, 1% BSA, 0.1% Tween 20, 250 mM NaCl overnight at 4°C. Samples were washed 3 times with 1xPBS + 0.3% Triton X-100 and incubated with appropriate secondary antibodies (from Jackson Immuno-Research Laboratories) for 2 hours at room temperature. Samples were mounted in 80% glycerol in a bridge formed by two cover slips to prevent the samples from being crushed while analyzed on the confocal microscope (Leica TCS SP5, 63X magnification oil immersion lens).

### Xbp1 RNA cleavage assay

T7 RNA was generated from pOT2-Xbp1, containing the ORF and UTR of *Drosophila* Xbp1. 1 μg of RNA was digested at room temperature by 1 μg of human IRE1α KR recombinant protein (~0.8 μM) for 45 min in RNA cleavage buffer (HEPES pH7.5 20 mM; K acetate 50 mM; Mg acetate 1 mM; TritonX-100 0.05% (v/v)). The total volume of the reaction is 25 μl. The digestion was then complemented by an equal volume of formamide and heated up at 70°C for 10 min to denature the RNA. The mixture was immediately placed on ice for 5 min, and then 20 μl was run on 3% agarose gel at 160V for 1 hour at 4°C. The PUM proteins were incubated with the RNA for 40 min on ice prior to RNA digestion.

### Statistical Methods

All panels data are represented as mean ± SEM, from at least three independent biological replicates experiments. All statistical comparisons for significance between control and experimental groups was calculated using a significance cut off p < 0.05. and denoted by *p< 0.05,**p< 0.01, and ***p< 0.001, based on two-tailed unpaired t-Student’s test or one-way ANOVA followed by an appropriate post-hoc analysis. Statistical analyses were performed using GraphPad Prism 8 (GraphPad Software, Inc) and online resources (https://astatsa.com/OneWay_Anova_with_TukeyHSD).

## AUTHOR CONTRIBUTIONS

FC designed the study, performed the experiments, analyzed data and wrote the manuscript. CS constructed reagents for the Phostag assays and contributed to the RNA IP assays. ALT conducted the Xbp1 mRNA in vitro cleavage assays. SM provided the purified versions of hPUM1. PD performed the IF staining in Fig.2. PD and AA designed and supervised the study, analyzed data and wrote the manuscript. All authors read and edited the manuscript.

## ACKNOWLEDGMENTS

We thank Aaron Goldstrohm for plasmids. We thank Yuh Nung Jan, Stefan Luschnig and André Gerber for Pumilio fly stocks. We thank David Ron and Heather Harding for providing the purified hIRE1α KEN, 4μ8C and helpful discussions. This work received funding by grants from the Fundação para a Ciencia e Tecnologia (PTDC/BIA-BCM/105217/2008; PTDC/SAU-OBD/104399/2008; PTDC/BEX-BCM/1217/2012; FCT-ANR/ NEU-NMC/0006/2013; PTDC/NEU-NMC/2459/2014; SFRH/BPD/93893/2013 and DL57/2016 to FC and SFRH/BD/ 130817/2017 to CS). The project leading to these results has also received funding from ‘la Caixa’ Foundation (ID 100010434), under the agreement <LCF/PR/HR17/52150018>.

---

**Figure S1.**
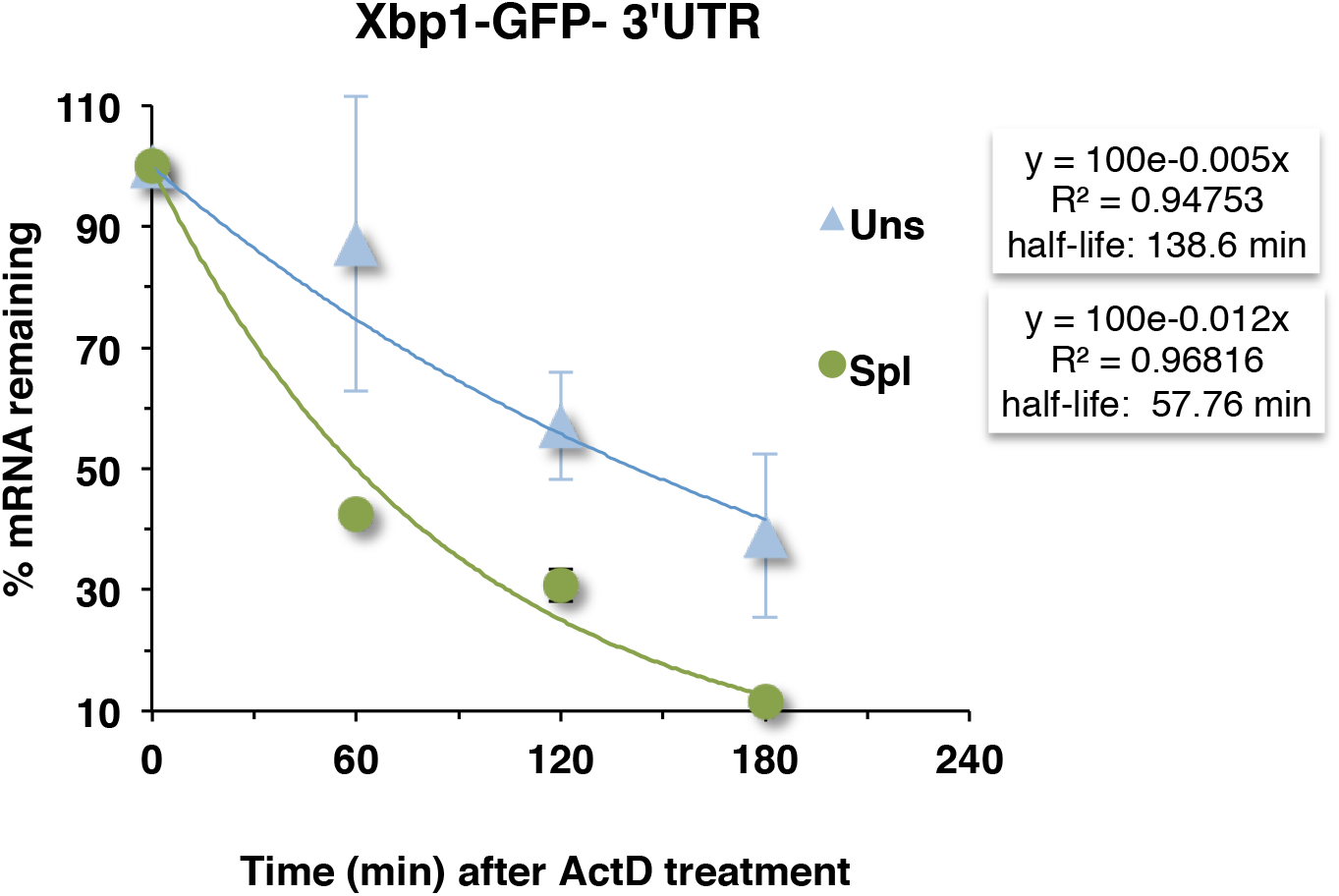
Xbp1 3’UTR ^Unspliced^ and 3’UTR^Spliced^ differ in their stability effect. Stability of the GFP reporters fused with Xbp1 3’UTR^spliced^ or 3’UTR^unspliced^ was assessed by quantitative RT-PCR (qRT-PCR), using primers specific for GFP and rp49 mRNAs (control). The levels of mRNA reporter were normalized to those of rp49 mRNA, and averages and standard deviations from three independent experiments are plotted. 3’UTR^unspliced^ shows a higher stability effect of the reporter half-life (2 fold stabilization) relatively to the spliced form.

**Figure S2.**
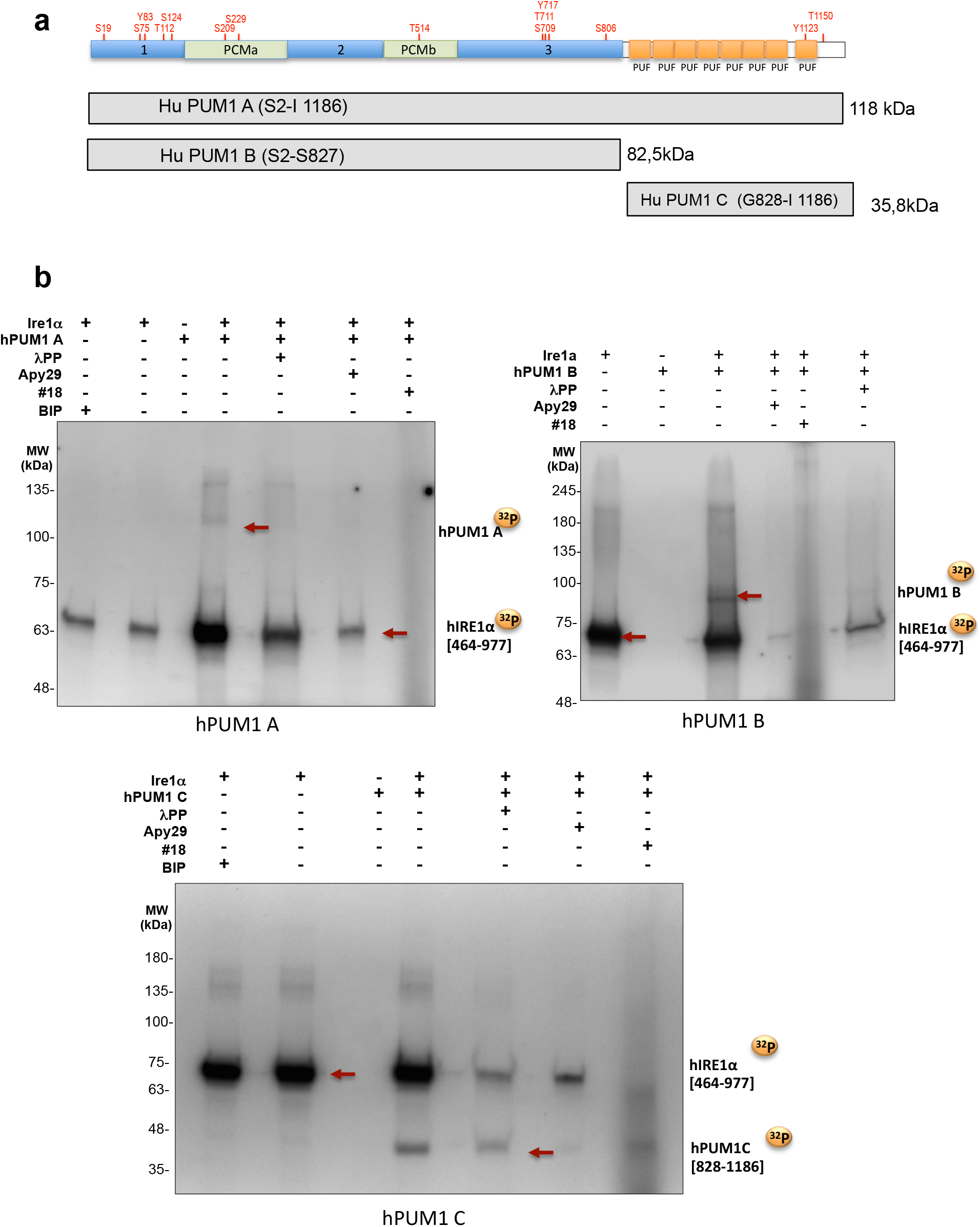
hIRE1 KR domain phosphorylates hPUM1 in vitro. **a)** Schematic representation of domains and putative phosphorylation sites present in Human PUM1. **b)** in vitro radioactive kinase assay with γ-[^32^P]ATP of purified hPUM1 forms incubated with hIRE1 KR domain. Specificity of phosphorylation was monitored by treatment with lambdaPP or incubation with specific inhibitors of IRE1 kinase activity (Apy 29 and #18). Arrows express hPUM1 phosphorylated forms corresponding to the expected MW of each purified protein. Treatment with kinase inhibitors decreased the amount of detected radioactive protein bands.

**Figure S3.**
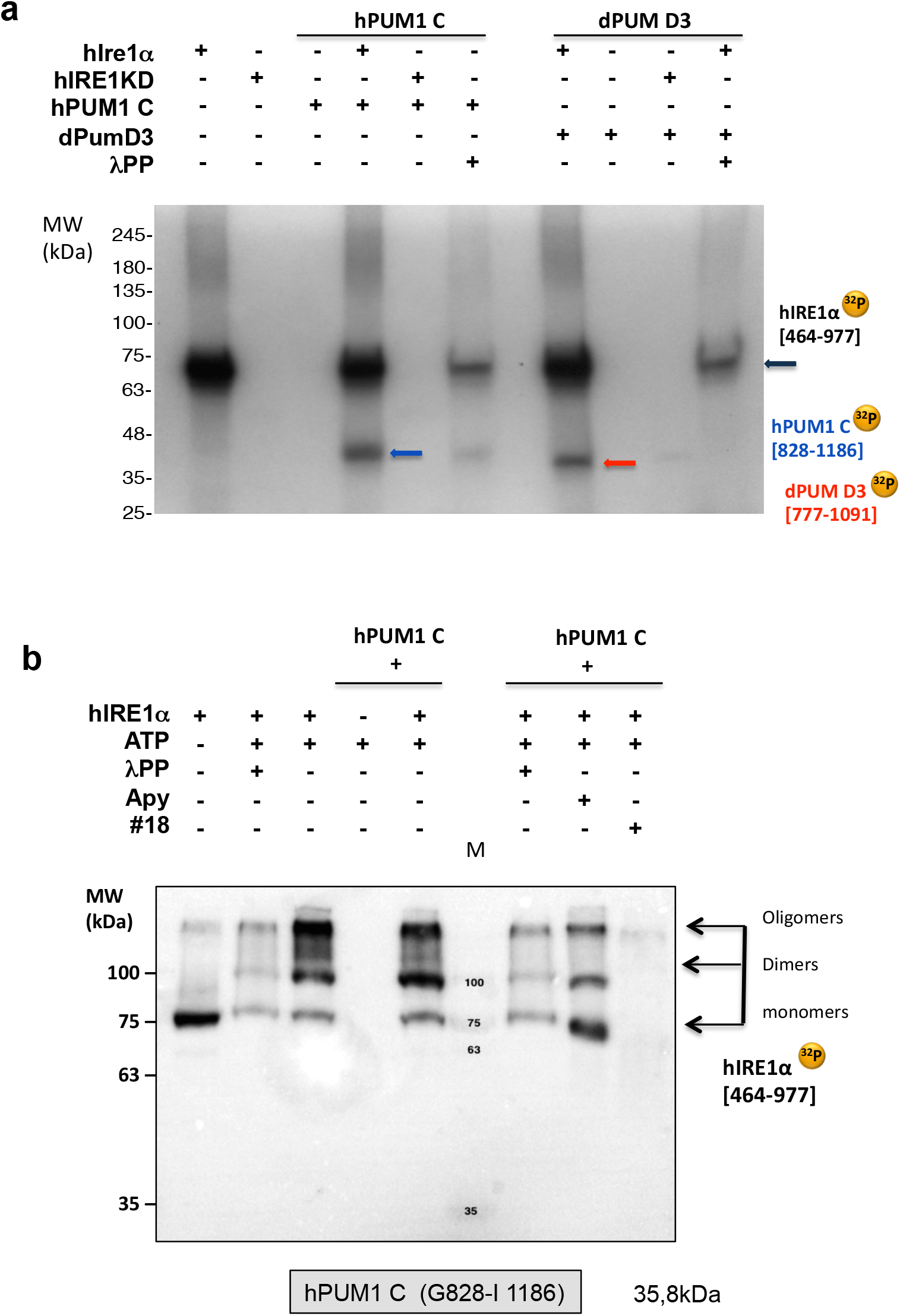
Validation of in vitro phosphorylation hPUM1-C and dPUM-D3. **a)** In vitro phosphorylation hPUM1-C domain and dPUM-D3 domain by hIRE1 KR using the control kinase-dead Ire1 (hIRE1KD, D688N). Assays were conducted as described before. Arrows denote phosphorylated forms of hPUM-C (Blue arrow) and dPUM-D3 purified proteins (red arrow). Autophosphorylation of hIRE KR is denoted by a black arrow. **b)** Phostag immunoblot of phosphorylation of hPUM-C with an antibody specific for phosphorylated IRE1(anti-rabbit Phospho-Ire1). Specificity of phosphorylation was monitored by treatment with λPP or incubation with specific inhibitors of IRE1 kinase activity (Apy 29 and compound #18) and a reaction lacking ATP (first lane). The phosphorylation state of hIRE1 KR is decreased in control reactions as expected comparatively to the reactions containing hIRE1+ATP or hIRE1 + hPUM-C + ATP. An additional control was made by conducting a kinase assay of hPUM-C without incubation with hIRE1 KR domain. M – Protein molecular weight marker lane.

